# Multiple extracellular polymeric substances pathways expressed by *Accumulibacter* and the flanking community during aerobic granule formation and after influent modification

**DOI:** 10.1101/2024.09.04.611157

**Authors:** Laëtitia Cardona, Jaspreet Singh Saini, Pilar Natalia Rodilla Ramírez, Aline Adler, Christof Holliger

**Author notes:** **Correspondence:** Laëtitia Cardona.

## Abstract

Aerobic granular sludge is a biological wastewater treatment process in which a microbial community forms a granular biofilm. The role of *Candidatus* Accumulibacter in the production of a biofilm matrix composed of extracellular polymeric substances was studied in a sequencing batch reactor enriched with polyphosphate-accumulating organisms. The metabolisms of the microbial populations were investigated using de novo metatranscriptomics analysis. Finally, the effect of decreasing the influent phosphate concentration was investigated.

A few weeks after the reactor start-up, the microbial community was dominated by *Accumulibacter*. Up to nine species were active in parallel. However, the most active species differed according to sampling time. Reducing the phosphate concentration led to a dominance of the glycogen-accumulating organism *Propionivibrio*, with some *Accumulibacter* species still abundant. *De novo* metatranscriptomic analysis indicated a high diversity of potential extracellular substances produced mainly by *Azonexus, Accumulibacter, Candidatus* Contendobacter, and *Propionivibrio*. Moreover, the results suggest that *Azonexus, Contendobacter* and *Propionivibrio* recycle the neuraminic acid produced by *Accumulibacter*. Changes in the microbial community did not cause the granules to disintegrate, indicating that a *Propionivibrio*-dominated community can maintain stable granules.

## 1 Introduction

Aerobic granular sludge (AGS) is a wastewater treatment process in which a microbial community grows in a self-supported granular biofilm. Enhanced biological phosphorus removal (EBPR) with this type of biomass can be achieved by operating a sequencing batch reactor (SBR) and alternating anaerobic feeding and aerobic starvation phases. This favours phosphate-accumulating organisms (PAO) responsible for EBPR, as well as glycogen-accumulating organisms (GAO). During the anaerobic feeding phase, these microbes take up the carbon source and store it intracellularly as polyhydroxyalkanoate (PHA). Simultaneously, PAO consumes intracellular polyphosphate and glycogen to provide energy and reducing equivalents, leading to the release of inorganic phosphate into the bulk liquid, whereas GAO relies only on glycogen. During the aerobic starvation phase, PHA is used to grow and replenish the reserves of polyphosphate (PAO only) and glycogen (both) (Nielsen et al., 2019).

AGS process is formed by a matrix of extracellular polymeric substances (EPS) that are synthesised and excreted by the microbial community. EPS comprise polysaccharides, proteins, nucleic acids and other molecules (Flemming et al., 2023). Destabilisation of the EPS matrix can result in a decrease in the fast settleability of the biomass, leading to the washout of the microbial community and a reduction in nutrient removal performance. Despite the importance of EPS in the AGS process, there is a scarcity of information regarding the microbial producers, regulators, EPS composition, and dynamics of these substances, particularly during granulation. Previous efforts to identify the EPS producer relied on isolation (Nouha et al., 2016) and metagenomic studies. In a recent study, Dueholm et al. (2023) compared the genetic potential of activated sludge bacteria to produce EPS based on their metagenome-resolved genomes. They identified specific operons in some of the most common microorganisms in AGS, such as poly-N-acetylglucosamine in *Candidatus* Accumulibacter (a PAO, referred to as *Accumulibacter*) and the Pel operon in *Propionivibrio* (GAO). A few studies conducted on SBR have shown that certain types of EPS are associated with enriched microbial populations. For example, exopolysaccharide ‘granulan’ has been extracted from *Candidatus* Competibacter-enriched biomass (Seviour et al., 2011), while complex glycoconjugates were identified in granules dominated by ammonium-oxidising bacteria (Lin et al., 2018). However, few studies have directly focused on EPS producers during AGS formation while identifying which microbes are producing specific types of EPS under certain condition is a pre-requisite in order to obtain a stable AGS process or an EPS production industrially viable, as they can be transformed into renewable bioresources (Karakas et al., 2020).

In this present work, we explore the use of metatranscriptomics analysis to dissect the role of an AGS microbial community during granulation, particularly regarding EPS production. Additionally, the influence of phosphate concentration on granule stability, microbial community activity, and gene transcription was assessed by decreasing the initially high phosphate concentration during the last eighty days of reactor operation.

## 2 Material and Methods

### 2.1 Reactor set-up

The experiment was carried out in a bubble column SBR of 6.2 cm diameter and 2.4 L working volume in fill-draw mode. The temperature was regulated at 18°C +/-1°C by recirculating water in the double-wall reactor. The pH was maintained at 7.5 +/-1 by regulating the injection of 1mM HCl or 1 mM NaOH with an ISFET probe (Endress+Hauser, Switzerland) using PID control. The pO2 and conductivity were monitored using two ISFET probes (Endress+Hauser).

A typical cycle corresponded to 5 minutes of sparging nitrogen gas to ensure anaerobic condition, 60 minutes feeding under anaerobic mixing condition, 45 minutes anaerobic phase to ensure total carbon consumption, 150 minutes aerobic phase by sparging compressed air, 15 minutes of sparging nitrogen to return to anaerobic condition, 5 minutes of settling and 5 minutes of withdrawal of half of the reactor working volume. The hydraulic retention time was set at 9.5 h and the solid retention time was set at 21 days by sampling at the end of the aerobic phase.

The durations of the different phases of the cycle were adapted across the experiment to ensure the optimal functioning of the reactor.

### 2.2 Inoculum and media

The sludge used as inoculum was collected in the anaerobic tank of ARA Thunersee (Thun, Switzerland), an activated sludge wastewater treatment plant that performs biological phosphorous removal. After transport (2-3h), the sludge was centrifuged 10 minutes, at 4°C and 5000 x g to concentrate the biomass. Then, the sludge was homogenised with a glass homogeniser with a distance between the pestle and the tube between 0.15 and 0.25 mm (50 cm3, Carl Roth, Germany). The total and volatile solid compositions were measured, and the sludge was stored at 4°C for a maximum of two days.

The reactor influent was created by mixing two 8.89 times concentrated solutions of C and NP and Milli-Q water. Concentrated solution C contained 5.67 g/L C_2_H_3_O_2_Na-3H_2_O, 2.28 g/L C_3_H_5_O_2_Na, 0.889 g/L MgSO_4_-7H_2_O, 2.2 g/L MgCl_2_-6H_2_O and 0.4 g/L CaCl_2_-H_2_O. Concentrated solution NP contained 1.671 g/L K_2_HPO_4_, 0.649 g/L KH_2_PO_4_, 0.048 g/L C_4_H_8_N_2_S to inhibit the nitrification and 50 mL of trace elements solution composed of 16.22 g/L C_10_H_14_N_2_Na_2_O_8_-H_2_O, 0.44 g/L ZnSO_4_-7H_2_O, 1.012 g/L MnCl_2_-4H_2_O, 7.049 g/L (NH_4_)_2_Fe(SO_4_)_2_-6H_2_O, 0.328 g/L (NH_4_)_6_Mo_7_O_24_-4H_2_O, 0.315 g/L CuSO_4_-5H_2_O and 0.322 g/L CoCl_2_-6H2O. Both solutions were autoclaved in 10 L glass bottles. Before use, 250 mL of bicarbonate solution composed of 0.933 g/L NH_4_HCO_3_ and 0.533 g/L KHCO_3_ was added to the NP solution to reach a final volume of 10 L. At each cycle, 120 mL of concentrated solutions C and NP were mixed with 960 mL of distilled water to feed the reactor and achieve a final chemical oxygen demand (COD) concentration of 300 mgO_2_/L in the SBR.

### 2.3 Description of the experiment

The reactor was inoculated with the concentrated and homogenised activated sludge to obtain a final concentration of 5 gMLSS/L. To limit the risk of carbon leakage during the aerobic phase, the COD concentration was increased stepwise from 50 to 300 mgO_2_/L within 15 days. Moreover, the anaerobic phase was decreased based on COD measurements from 150 minutes at the beginning of the experiment to 45 minutes at the end. The settling time was decreased from 50 to 5 minutes when granule formation and fast biomass settling was observed.

The SBR was operated for 183 days. After 103 days of operation, the COD/P ratio was increased from 12 to 200 by decreasing the phosphate concentration in the influent. Because the phosphate concentration released at the end of the anaerobic phase remained high, two consecutive cycles were operated differently. These conditioning cycles were modified as follows: at the end of the anaerobic phase, the reactor was stopped to allow the biomass to settle, and half of the working volume was removed before the start of the aerobic phase to reduce the formation of polyphosphate reserves. This operation was repeated one week later.

### 2.4 Nutrient removal performance monitoring

Nutrient removal performance of the reactors was measured weekly. Samples of 50 mL were taken from the SBR at the end of the anaerobic and aerobic phases, centrifuged 5 minutes at room temperature at 4200 x g and the supernatant was filtered (0.45µm). A sample of the synthetic reactor influent was also collected and filtered (0.45µm). The samples were stored at 4°C until further analysis. The concentrations of the anions (P-PO4^3-^, N-NO3^-^, and N-NO2^-^) were measured using ionic chromatography (IC, ICS-90, IonPacAS14A column) with an electrical conductivity detector (Dionex, Switzerland). The Chemical Oxygen Demand was measured by spectrophotometry using two different kits: LCK514 (100-2000 mgO2/L) and LCK 314 (15-150 mgO2/L) (Hach, USA), measured on a spectrophotometre DR 3900 (Hach, USA).

The total and volatile solids were determined in the sludge obtained by centrifuging 100 mL of mixed liquor reactor sample taken at the end of the aerobic phase. The mass of the dried pellet after 12 h of drying at 105°C yielded the total solids, and the mass loss after 2 h of calcination at 550°C resulted in volatile solids.

### 2.5 AGS granule size and morphology

Granule formation was followed by capturing a picture of the biomass using a camera from a Dino-Lite Edge Digital Microscope and the DinoCapture 2.0 software (AnMo Electronics Corporation, Taiwan). The mean particle size distribution was measured weekly using an LS13 320 Series particle size analyser (Beckman Coulter, Germany) connected to a universal liquid module. The particle range goes from 0.37 to 2000μm and the following parameters were used for the measurements: obscuration 12%, optical model Fraunhofer, run length of 60 seconds, pump flow of 20%.

### 2.6 16S rRNA gene amplicon sequencing and analysis

The sampling for the 16S rRNA gene amplicon sequencing, the DNA extraction and 16S rRNA gene amplicon sequencing (protocol no.2) were performed as described by Adler and Holliger (2020). Briefly, the samples were taken every week at the end of the anaerobic phase, washed with PBS 1X solution, homogenized with a glass pestle and stored in -20°C until use. The extraction was performed using the automated robot 16 DNA purification system (Maxwell, Promega Corporation, Switzerland) after enzymatic lysis (lysozyme, 1h at 37°C). The bacterial 16S rRNA gene hypervariable regions V1-V2 were amplified using the universal primers 27F and 338R and High-Fidelity Q5 polymerase (High-fidelity 2x Master Mix, Biolabs Inc., USA). A secondary indexing PCR was performed on normalised samples at 5 ng/μL using TG Nextera XT Index kit v2 Set B (#FC-131-2002, Illumina, USA). The products were purified using Agencourt AMPure XP magnetic beads and quantified using Qubit dsDNA HS. Pooling of the normalised samples was performed at 10 nM, and sequencing was performed by the Lausanne Genomic Technologies Facility (University of Lausanne, Switzerland) on an Illumina MiSeq v2 in paired-end mode (2 × 250).

Adapter sequences were removed from the reads using cutadapt v3.5 (Martin, 2011) and default parameters after removing Ns from the sequences using DADA2 v1.32.0 (Callahan et al., 2016) with filterAndTrim function. Then, reads were quality-filtered and trimmed using the filterAndTrim function (truncLen=c(225,225), maxEE=c(2,2), truncQ=2, maxN=0). Error rates were estimated for both forward and reverse sequences using learnErrors (nbases = 1e10, randomize = TRUE). Sequences were dereplicated and inferred using the derepFastq and dada (pool = “pseudo”) functions. Paired-end sequences were merged using the mergePairs function before removing the chimera with removeBimeraDenovo (method = “consensus”). Finally, the taxonomy was assigned using the assignTaxonomy function with the MIDAS 5.3 database (Dueholm et al., 2024). In total of 7138 ASVs were obtained.

Low abundant ASVs were filtered out if there were not a minimum of 15 reads in at least 20 % of the samples (538 ASVs out of 7138). The filtered ASVs were then agglomerated to the genus level using the glom_tax function of the phyloseq package. After filtering and agglomeration, 234 genera remained in total. The full taxonomic information and counts of the processed genera are provided in Appendix B.

### 2.7 Metatranscriptomics

For the metatranscriptomic analysis, four biomass samples were collected on days 13, 26, 103, and 182 to capture the microbial activity at different stages of granule formation. An aliquot of 15 ml of mixed liquor was sampled after 15 minutes of feeding and after 15 minutes of aeration on 3 consecutive cycles. The samples were quickly placed on ice and centrifuged for 1 min at 4°C and 4200 x g. The supernatant was filtered (0.45µm) to measure the COD and P. The pellet was resuspended in 2 volumes (g pellet:ml volume) of RNA protect Tissue (Qiagen, Germany) and homogenised by passing 3 times through a needle (26G). After an incubation at room temperature (RT) for 5 minutes, the sample was centrifuged 5 minutes at RT and 5000 x g and the supernatant was discarded. The pellet was snap-frozen in liquid nitrogen and stored at -80°C until RNA extraction.

For RNA extraction, the samples were thawed on ice, resuspended in 0.5 mL of TRIzol (#15596-0026, Invitrogen, Fisher Scientific AG, Switzerland), and incubated for 5 minutes at RT. Then, 0.1 mL of chloroform 99+% was added, and the mixture was vortexed for 15 seconds and incubated 2 minutes at RT before being centrifuged 15 minutes at 15500 x g at 4°C. The upper portion was recovered and mixed with 400 µL of 100% ethanol. RNA was purified using an RNA purification kit (Direct-zol RNA Miniprep #R2050, Zymo Research, Germany) following the manufacturer’s recommendations, with the exception of centrifugation performed for 1 minute at 13000 x g. DNA was removed using a TURBO DNA-*free*™ kit (#AM1907, Thermo Fisher Scientific, Switzerland) following the manufacturer’s recommendations. RNA was purified using magnetic beads (Agencourt RNACleaner XP, #A63987, Beckman Coulter) by adding 76 µL magnetic beads to the extracted RNA. After mixing 10 times up and down, the samples were incubated at RT for 5 minutes. The samples were placed on the magnetic rack for 10 minutes before removing the supernatant followed by the addition of 200 µL of ethanol 70% and incubation of 30 secondes. The ethanol was removed, and the previous steps were repeated twice. After removing the ethanol, 32 µL of RNAse and DNase free water was added to the pellet out of the rack and resuspended 10 times by up and down. The samples were incubated for 1 minute before being returned to the rack for 1 minute. Finally, the supernatants were collected. Bacterial ribosomal RNA (rRNA) was removed using a QiaSeqFastSelect 5S/16S/23S kit (#335925, Qiagen) following the manufacturer’s recommendations on 1 µg of total RNA (protocol with TruSeq^®^ stranded library preparation) with the following modification: the first step of combined fragmentation and hybridisation was performed for 1 minute at 89°C. Libraries were then generated using the Truseq Stranded mRNA sample preparation kit (#20020594, Illumina, USA) and IDT for Illumina TruSeq RNA UD Indexes (#20022371, Illumina) following the reference guide #1000000040498 for the LS procedure without optional steps. For the clean-up amplified DNA step, the ratio of magnetic beads: PCR products was 0.7 and 20 µL of RSB was added to release the genetic material from the bead. The amplification was quantified with Qubit dsDNA HS Assay Kit (#Q32854, Life Technologies) and the quality was checked by electrophoresis using Agilent High Sensitivity DNA Kit (# 5067-4626, Agilent Technologies). The concentrations of the samples were normalised to 10 nM and pooled. Sequencing analysis was performed at the Lausanne Genomic Technologies Facility, University of Lausanne (Switzerland) on a NovaSeq 6000 in paired-end mode (2 × 150).

Before *de novo* metatranscriptomics analysis, the quality of the reads was evaluated using FastQC v0.11.9 (https://www.bioinformatics.babraham.ac.uk/projects/fastqc/) for the raw and at each step of the treatment. The results were summarised using multiqc v1.13 (Ewels et al., 2016). The reads were filtered and trimmed using BBDuk from BBMap v39.01 (https://jgi.doe.gov/data-and-tools/software-tools/bbtools/bb-tools-user-guide/bbmap-guide/) using the following parameters *ktrim=r, k=23, mink=11, hdist=1, tpe, tbo* for the adapter trimming steps and *qtrim=rl, trimq=20, minlen=50, maq=20, maxns=1* for the quality trimming and filtering. Ribosomal RNA was removed using sortMeRNA v4.3.6 (Kopylova et al., 2012) and the databases for silva-bac-16s-id90, silva-arc-16s-id95, silva-euk-18s-id95,silva-bac-23s-id98, silva-arc-23s-id98, silva-euk-28s-id98, rfam-5s-database-id98, rfam-5.8s-database-id98 from Silva and rfam. RNASpades (Bushmanova et al., 2019) was used to co-assemble the reads, and the results from the hard filtering from RNASpades were used for the following steps. The quality of the obtained assembly was evaluated using BUSCO 5.4.7 and the bacteria_Odb10.2020-03-06 (*-m transcriptome*) (Manni et al., 2021) (Complete: 65.3%[Single-copy: 18.5%, Duplicated: 46.8%], Fragmented: 7.3%, Missing: 27.4%, n:124) and bowtie2 v2.4.1 (Langmead and Salzberg, 2012) *(-k 20*) by mapping back the reads onto the co-assembly (95 % of reads mapped onto the assembly). The redundancy in the sequences was considered by clustering the redundant information using CD-HIT v4.8.1 (Li and Godzik, 2006) (*-c 0*.*95, -n 10, -d 0, -M 16000*). FeatureCount v2.0.1 (Liao et al., 2014) was used to summarise the count in a table using the following parameters *-O -M -C -B –fraction* (360881 annotated genes) and the results are provided in the supplementary file 2. The pipeline results are summarised in Appendix C. DRAM v1.4.6 (Shaffer et al., 2020) and eggNOG-mapper v2.1.11 (Cantalapiedra et al., 2021) were used for gene prediction and annotation. For the transcripts of interest highlighted in this paper, the amino sequences were blasted on NCBI using BLASTP to check the taxonomic assignment and gene description. If the percentage of identity was under 80%, the transcript was not considered for further analysis. Kaiju v1.9.9, and the non-redundant protein database (update 22 March 2022) (Menzel et al., 2016) was used to obtain taxonomic assignments at the genus level of the assemblies. Gene and taxonomic information are summarized in Appendix D. mOTUs v3 (Ruscheweyh et al., 2022) was used to obtain the expression profile at the species level after extending the database 2.8, metagenomes metagenome-assembled genomes obtained previously from the laboratory (Saini et al., 2024) and from other publications (Arumugam et al., 2021; Singleton et al., 2021; Ye et al., 2020), following the procedure described in the github page (https://github.com/motu-tool/mOTUs-extender).

### 2.8 Microbial community activity analysis

All the statistical analyses were done on R (v4.3.3) and RStudio (2023.12.1+402) and the graphics were obtained using ggplot2 (v3.5.1). Low count filtering (minimum 15 counts) was applied using *filterByExpr* from edgeR (v4.0.16) (Robinson et al., 2010) on the metatranscriptomics count results. After filtering, 184899 genes remained. The expression of each transcript was normalised using *normLibSizes* (method = “TMM”) from edgeR. The results of the raw, filtered, and normalised counts are provided in Appendix C.

The relative activity of the active members at the genus level was obtained by summing the cpm values, resulting from the TMM normalisation, of all the transcripts associated with each genus. Alpha diversity using the Shannon index was calculated for this genus-level dataset.

Genes involved in specific metabolic pathways were selected based on KO, COG, and PFAM identification numbers. The complete list of the genes of interest is provided in Appendix E. For these selected transcripts, the sum of the expression was calculated for each gene, genus, and sample. The final expression level was obtained by calculating the median of each gene considering the cycles in triplicate.

Differential gene expression (DGE) analysis was performed using edgeR to compare the anaerobic and aerobic phases for each time point. As only 15% of the genes were identified as differentially expressed, we focused on DGE analysis between time points which identified approximately 60% of the genes as differentially expressed. However, the phases were kept separate for the comparison of each time-point analysis; only the cycles were considered as replicates for each sample. Only the results for the feeding samples are presented in the main article, as both results for feeding and aerobic conditions were highly similar for the analyzed genes. The results for the aerobic samples are presented in Appendix A, Figures S2 and S3, respectively.

## 3 Results

### 3.1 Nutrient removal performances

In this study, the carbon removal efficiency was always higher than 90% after an adaptation phase of 15 days (Figure 1.A). As expected for typical PAO metabolism, the phosphate concentration increased during the anaerobic phase, reaching up to 250 mg/L, and then decreased to very low concentrations during the aerobic phase (Figure 1.B). When the influent phosphate concentration was drastically reduced (day 103), the value at the end of the anaerobic phase remained high. This was probably due to the polyphosphate reserves of PAO. After the conditioning cycles on days 120 and 127 where phosphate-rich bulk liquid was removed at the end of the anaerobic phase, the phosphate release progressively decreased to 1-15 mg/L.

**Figure 1.**
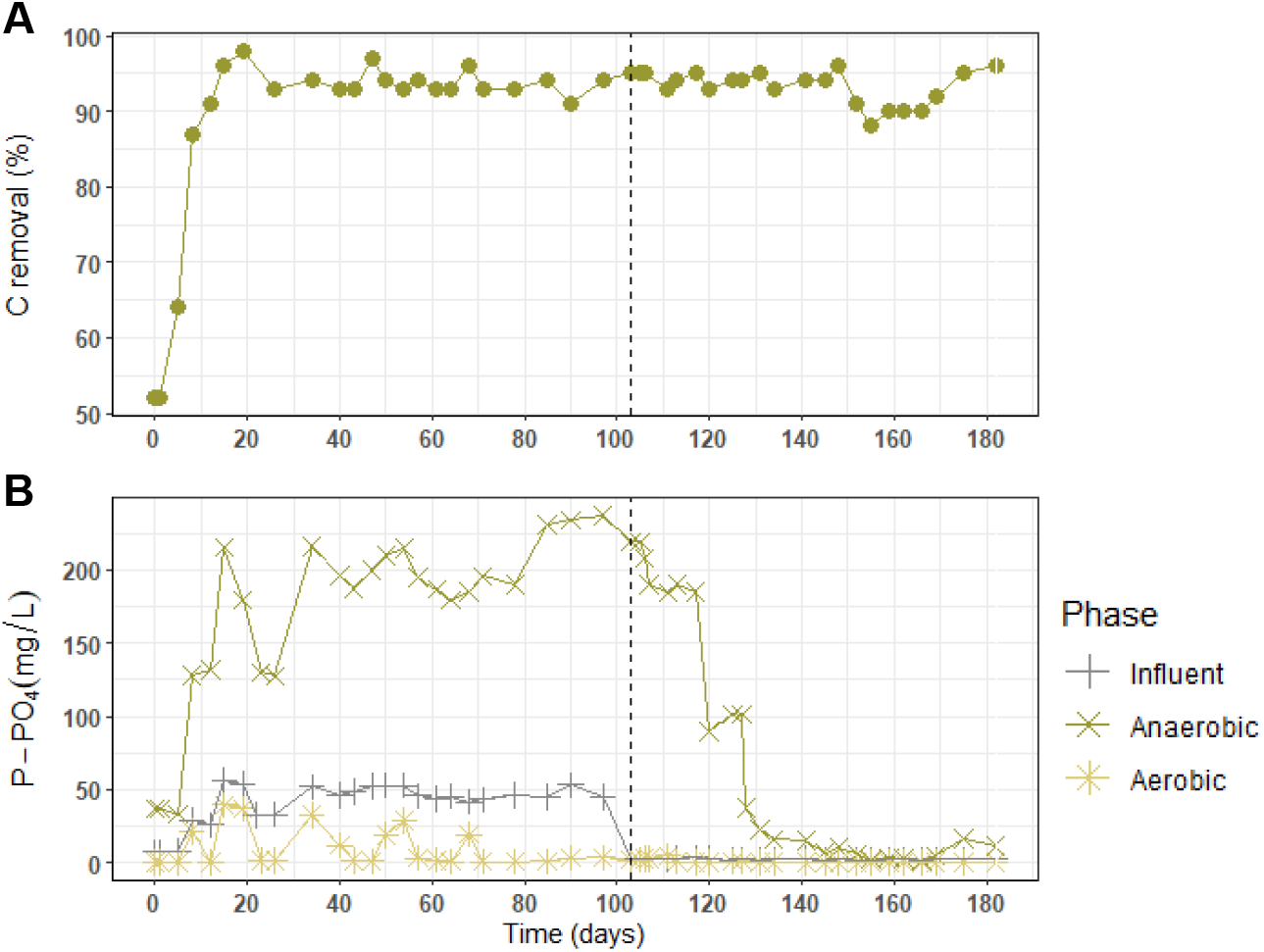
Nutrient removal and efficiency. A) Carbon removal efficiency during the anaerobic phase. B) Phosphate concentration at the different phases: influent, end of anaerobic and end of aerobic phase. Dashed line at day 103 indicates the change in the influent composition from COD/P of 12 to 200.

### 3.2 Granulation and microbial composition dynamics

The relative abundance of different populations of the microbial community is represented in the Figure 2. At the early beginning of the experiment, the microbial community was more diverse and mainly composed of *Candidatus* Phosphoritropha (formerly *Tetrasphaera* clade III), Nitrospira, a nitrite-oxidising organism, the metabolically versatile *Rhodobacter*, and many other microorganisms with a lower abundance. After a week of reactor operation, the relative abundance of these microorganisms decreased while the relative abundance of the PAO *Accumulibacter* (40 %), *Flavobacterium* (10%), the GAO *Contendobacter* (10%), Hyphomonadaceae UKL13-1(10%) and few other uncharacterised genera from the Flavobacteriaceae and Micavibrionales increased.

**Figure 2.**
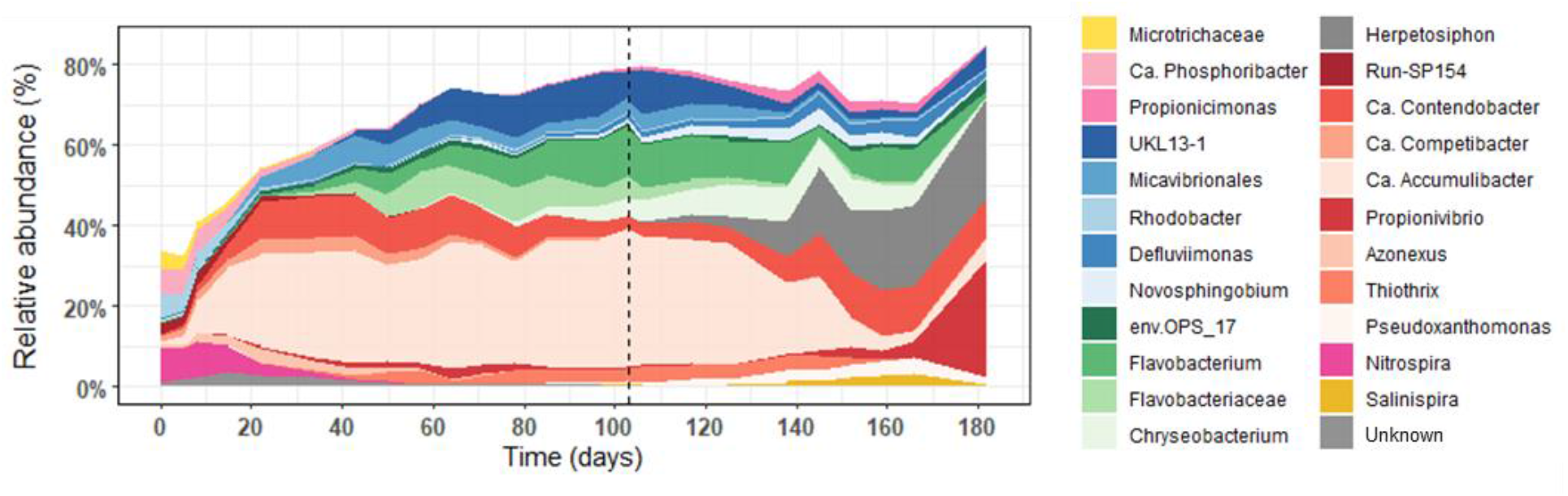
Relative abundance of the most abundant microorganisms. Relative abundance of the most abundant genera (minimum 3% in at least one sample) based on the 16S rRNA gene amplicon sequencing. The grey dashed line at day 103 indicates the change in the medium composition (from COD/P of 12 to COD/P of 200).

After the influent phosphate concentration was lowered at day 103, the microbial community composition slightly changed during the first 20 days. A progressive increase of the relative abundance of *Herpetosiphon* (25 %), *Salinispira* (3 %) and *Pseudoxanthomonas* (3 %) was observed. From day 125 onwards, the relative abundance of *Accumulibacter* progressively decreased to 5% while the one of *Propionivibrio* increased mainly at the end of the experiment to 28 %.

The aspect of the biomass changed across the experiment (Figure S1). From the initial flocs and little aggregates smaller than 200 µm (diameter), the biomass formed tiny smooth granules that grew to over 1000 µm in size. The aspect of the granules changed from smooth to grainy during the low-phosphate period.

### 3.3 Active members of the microbial community

In order to identify the active microbial community members at different stages of granulation, metatranscriptomics samples were taken at day 13 (mean particle size around 172 µm, COD/P ratio of 12), day 26 (300 µm, COD/P ratio of 12), day 103 (>1000 µm, COD/P ratio of 12) and day 182 (>1000 µm, COD/P ratio of 200).

When the granules were still small (days 13 and 26), *Accumulibacter* was the predominant active population accounting for 70% of the transcripts (Figure 3.A). When the granules were larger (day 103), *Accumulibacter* remained predominant but other active populations such as *Flavobacterium, Kaistella, Thiotrix* and *Azoarcus* emerged. The Shannon diversity index confirmed the increase of the active members diversity with increased granule size (Figure 3.B).

**Figure 3.**
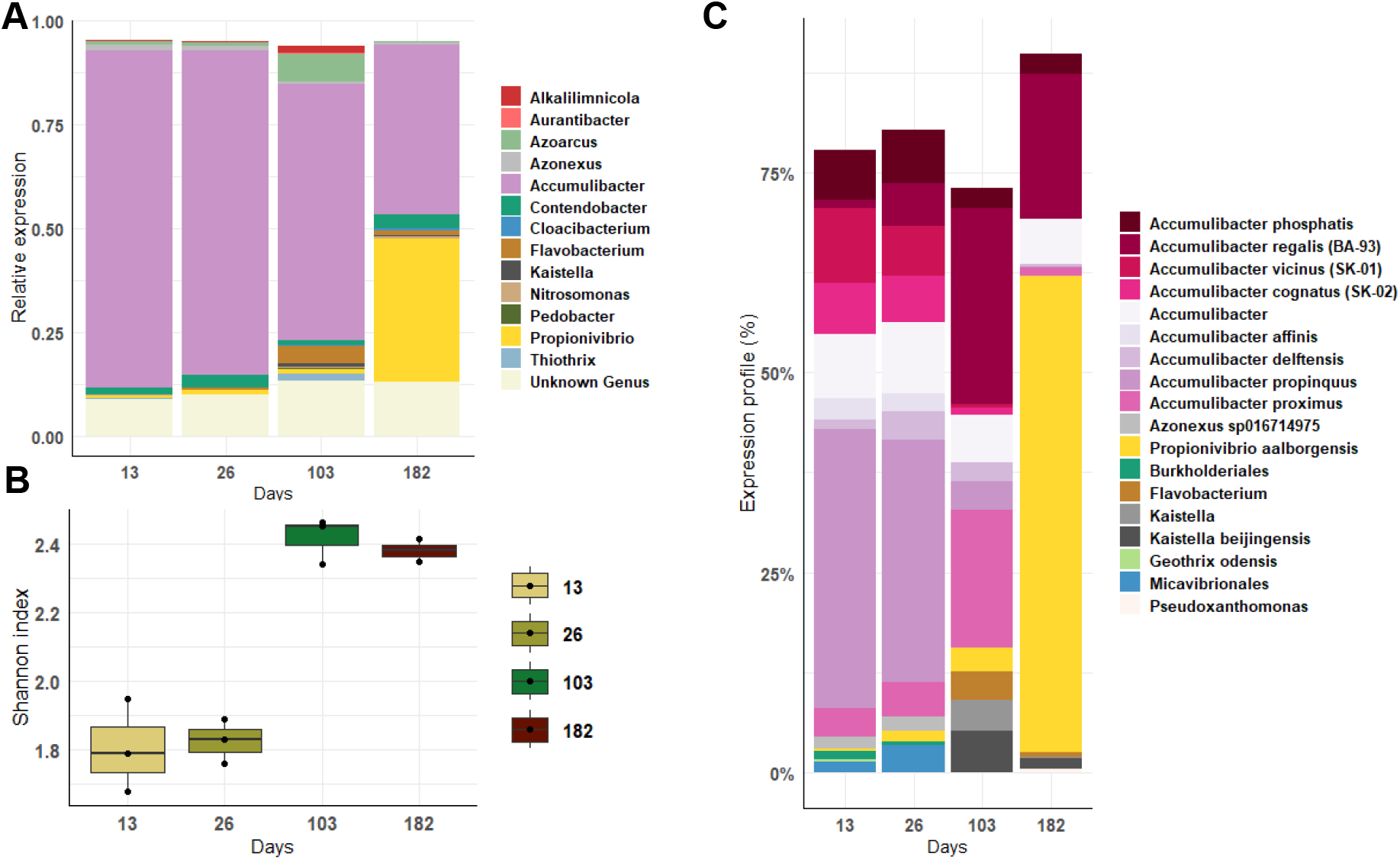
Active microbial population at different stages of granulation. A) Relative proportion of gene expression per genus from the de novo metatranscriptomics analysis. Only the 20 most abundant genera are represented. B) Shannon diversity index calculated using the level of expression of each genus. C) Expression profile at species level of the most abundant microorganisms using mOTUs (minimum of 3% in one sample).

At the end of the low-phosphate period (day 182), *Accumulibacter* and the GAO *Propionivibrio* were the dominant active populations. Although the conditions were favourable for the enrichment of GAO, only *Propionivibrio* was able to occupy the niche while the gene expression of *Candidatus* Contendobacter, another GAO detected in all the samples, remained low.

The co-occurrence of multiple active species of *Accumulibacter* was observed at the different stages of granulation (Figure 3.C). Interestingly, a shift in the main active ones was observed between day 26 and day 103. Whereas *Accumulibacter propinquus* was the most active species in small granules, *Accumulibacter regalis* BA-93 and *Accumulibacter proximus* predominated in large granules. Under the low-phosphate period, *Accumulibacter regalis* BA-93 remained the main active PAO.

### 3.4 Expression of metabolic pathways involved in EBPR

The microorganisms involved in metabolic pathways typical for the EBPR process were identified by analysing the expression of the genes related to acetate and propionate uptake and activation and the metabolisms of polyhydroxyalkanoate (PHA), glycogen, and polyphosphate (Figure 4). *Accumulibacter, Contendobacter*, and *Propionivibrio* were the main genera involved in these processes. *Candidatus* Competibacter (GAO) expressed few genes of the different metabolisms which could be due to its low abundance throughout the experiment.

**Figure 4.**
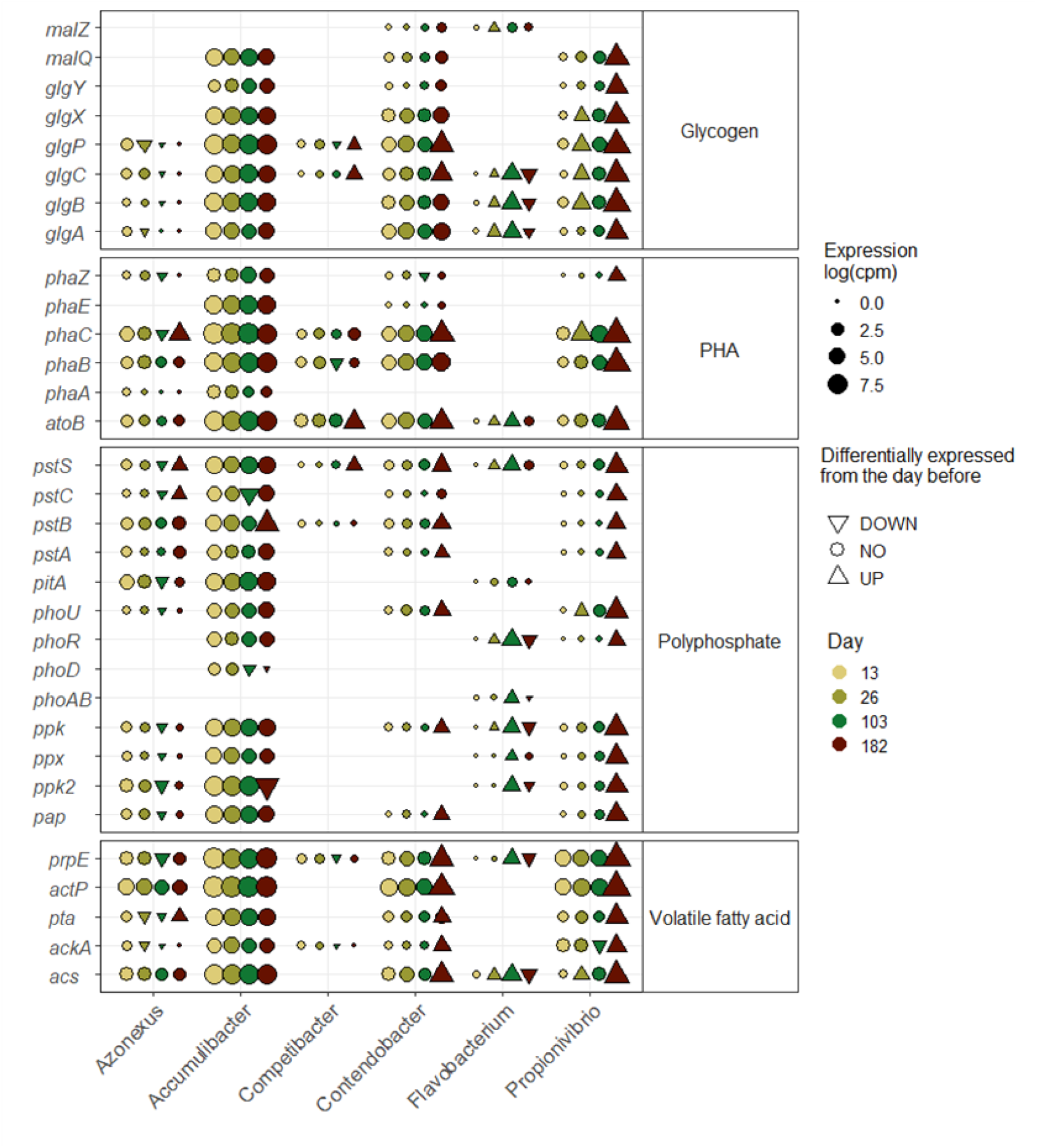
Expression of Enhanced Biological Phosphate Removal related genes at the feeding phase. Level of expression (log(cpm)) of genes per day for different genera. Differential gene expression analysis was done between two time points (26 versus 13, 103 versus 26 and 182 versus 103) and the significative differences (log-fold change > 2 and pvalue < 0.01) are represented by a triangle (up-pointing for up regulation and down-pointing triangle for down-regulation).

*Contendobacter* and *Propionivibrio* expressed polyphosphate-related genes except the low affinity transporter *pitA*. While the overall level of expression of *Contendobacter* did not increase along the experiment, genes related to the EBPR process of Propionivibrio had a very high level of expression at the end of the low-phosphate period.

In this experiment, *Azonexus* expressed genes related to acetate and propionate uptake (*actP* and *prpE* respectively) and activation (*acs, ackA* and *pta*), high and low phosphate affinity transporters (*pstABCS* and *pitA* respectively), polyphosphate kinases (*ppk* and *pap*), genes involved in PHA formation (*phaA, B* and *C*) and degradation (*phaZ*) and in glycogen branching (*glgABC*) and debranching (*glgP*).

Besides known PAO and GAO genera, *Flavobacterium* also expressed genes from EBPR pathways *i*.*e*. polyphosphate metabolism, glycogen branching and propionate transport. The level of expression for the different genes in *Flavobacterium* increased over time while the phosphate concentration was high. However, when the influent phosphate was low, the level of expression decreased.

### 3.5 Expression of extracellular polymeric substances

At all stages, *Accumulibacter* expressed genes from nonulosonic acids (N-acetylneuraminic acid, legionaminic acid (leg) and pseudaminic acid (pse)), cellulose, alginate and poly-N-acetylglucosamine (PNAG) pathways (Figure 5). Interestingly, genes for cellulose, PNAG and pse pathways were down-regulated with the increase of the granule size, while the change in the medium composition did not seem to influence their expression. In contrast, genes of the leg pathway were up-regulated. Regarding the nonulosonic acids, nonulosonic synthase (*neuB*) transcription decreased with time while the expression of neuraminic acid binding protein (*siaP*), permease (*dctM*) and tripartite ATP-independent periplasmic transporter proteins (*dctQ*), that enable the import of neuraminic acid, remained stable. Similarly, *Propionivibrio, Azonexus* and *Contendobacter* also expressed the genes related to the recycling of neuraminic acids but not *neuB*.

**Figure 5.**
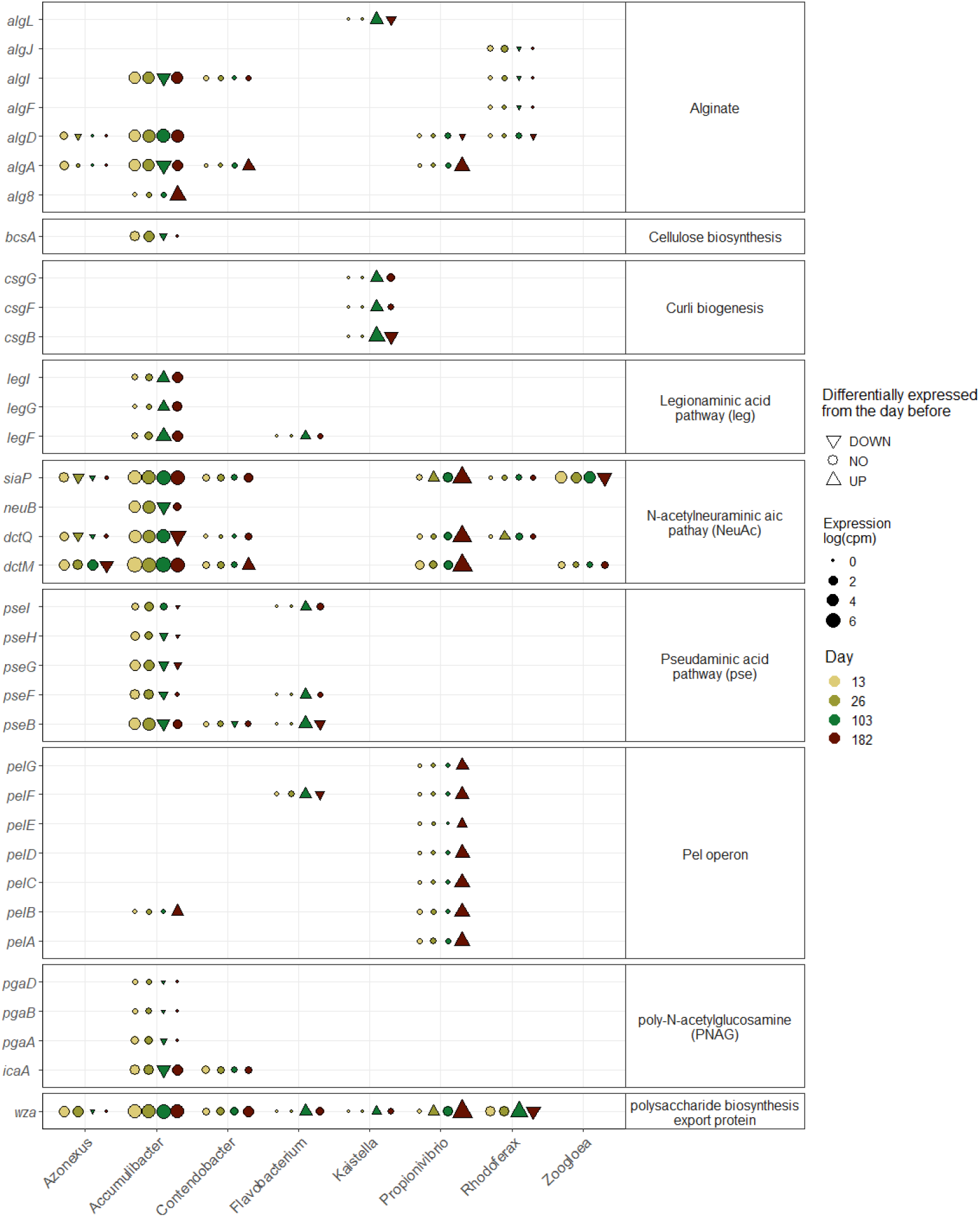
Expression of biofilm related genes at the feeding phase. Level of expression (log(cpm)) of genes per day for different genera. Differential gene expression analysis was done between two time points (26 versus 13, 103 versus 26 and 182 versus 103) and the significative differences (log-fold change > 2 and pvalue < 0.01) are represented by a triangle (up-pointing for up regulation and down-pointing triangle for down-regulation).

Different populations seemed to be involved in the production of other EPS at different stages of the experiment. Most major populations expressed the polysaccharide biosynthesis export protein *wza*, common in gram-negative bacteria (Ford et al., 2009). *Propionivibrio* expressed the pel cluster genes, *Rhodoferax* and *Azonexus* transcribed some genes from the alginate pathway, mostly at the beginning of the experiment, and expression of genes from the curli biogenesis pathway of *Kaistella* were detected.

Finally, *Zoogloea* only expressed two genes from the neuraminic acid pathway. However, in this study, the overall expression of *Zoogloea* was lower than 0.01%, and if it played a role during the granulation, it was not captured in the metatranscriptome.

## 4 Discussion

### 4.1 Microbial population dynamics during aerobic granulation

Phosphate and COD dynamics in the SBR suggested PAO metabolism. During the anaerobic phase, the carbon sources acetate and propionate were consumed, and phosphate was released, while in the aerobic phase, phosphate was consumed. Metatranscriptomics confirmed high activity of *Accumulibacter*, a widely present and abundant PAO (Petriglieri et al., 2022). Concurrently, round granules increased in size. Various *Accumulibacter* species were active at different granulation stages, with species abundance changing as granules grew. This co-occurrence has been noted in full-scale plants (Flowers et al., 2013) using clade-specific qPCR primers and in laboratory SBRs (Adler et al., 2022) using metagenomics. Both studies observed species dynamics related to temperature or influent composition, respectively. Leventhal et al. (2018) found that different *Accumulibacter* species co-occur in reactors but occupy distinct niches within granules. In this study, species-level expression profiles confirmed the presence and variation of *Accumulibacter* species, likely due to granule size changes, as no other parameter varied.

Microorganisms were active alongside *Accumulibacter*, albeit in lower abundance. The Shannon index showed increased diversity among active members with larger granule size. Similarly, Tan et al. (2014) noted this phenomenon using complementary DNA and suggested that micro-niches within larger granules support a wider variety of bacterial populations. The community included GAO *Contendobacter* and *Propionivibrio*, which, like PAO, can take up acetate and propionate during the anaerobic feeding phase. *Azonexus*, previously *Dechloromonas*, was active mainly during the first two time points. Metagenomics identified some *Azonexus* species, such as *A. phosphorivorans* and *A. phosphoritropha*, as PAO, with genes related to polyphosphate, glycogen, and PHA accumulation (Petriglieri et al., 2021). In this study, *Azonexus* expressed genes for phosphate transport and accumulation, PHA metabolism, and glycogen metabolism, indicating *Azonexus* (*Azonexus* sp016714975 in GTDB-tk, *Candidatus* Dechloromonas phosphorivorans in NCBI) was involved in the EBPR process.

It is noteworthy that *Flavobacterium* exhibited increased activity during granulation, and expressed genes related to the EBPR metabolism, particularly those involved in phosphate metabolism, volatile fatty acid activation, and glycogen branching. However, no genes from the PHA metabolism were identified as being expressed. Previous genomics studies of *Flavobacterium* sp. have identified genes such as *pitA, pstSCAB, ppk*, and *ppx*, which are thought to play a role in phosphate accumulation in the wetland environment from which this bacterium was isolated (Choi et al., 2023). Additionally, the presence of *glgABC* has been reported in psychrophilic species and was proposed as a means to resist low temperatures (Liu et al., 2019). Despite their prevalence in activated sludge (Dueholm et al., 2022), the role of *Flavobacterium* in AGS remains unclear. Further investigation is warranted to elucidate the significance of *Flavobacterium* in AGS.

### 4.2 *Accumulibacter* and *Propionivibrio* as main active population under low-phosphate condition

The phosphate concentration in the influent was decreased after 103 days to study the effect of low-phosphate condition on the microbial community activity and the granule stability.

When significantly reducing the influent phosphate concentration, the anaerobic phase’s phosphate levels remained high, and the microbial community, dominated by *Accumulibacter*, did not change, likely due to PAO’s polyphosphate reserves. The phosphate concentration only decreased after two anaerobic post-drainage cycles to remove excess phosphate. During this period, the microbial community composition began to change. By the experiment’s end, the absence of phosphate release/uptake and efficient carbon removal during the anaerobic phase indicated typical GAO metabolism (Acevedo et al., 2012). Microbial community composition and activity analyses supported this hypothesis, with *Propionivibrio* becoming abundant and active mainly at the experiment’s end. Interestingly, the GAO *Contendobacter*, present from the beginning, maintained constant relative abundance and activity, suggesting it was less competitive than Propionivibrio in optimal conditions. This may be due to operational conditions, presence of propionate, pH, temperature, or SBR phase operations being sub-optimal for *Contendobacter* development.

Based on the 16S rRNA gene amplicon sequencing results, relative abundance of *Herpetosiphon* increased after the change in the influent composition. *Herpetosiphon* are ubiquitous, filamentous bacteria and have already been identified in PAO-enriched reactor (Weissbrodt et al., 2013).They also have the ability to prey on other microorganisms (Livingstone et al., 2018). However, from the metatranscriptomics results, *Herpetosiphon* was barely active, the role of this genus was then impossible to display. The discrepancy between the two analyses can be explained by the bias of each method (different extractions, library preparation, sequencing depth…) but could also be part of the difference between being present and active.

*Accumulibacter*, considered as classical PAO, can shift to GAO metabolism under low phosphate conditions. Indications have been obtained that both *Accumulibacter* types I and II can alter their metabolism, with type I being less competitive due to a lower acetate uptake rate under polyphosphate-depleted conditions (Welles et al., 2015). The division into two types is based on the polyphosphate kinase gene (ppk1) and supported by genome-resolved phylogenies (Petriglieri et al., 2022). The current study’s mOTUs profiling revealed two *Accumulibacter* types I (*A. regalis, A. delftensis*) and II (*A. propinquus, A. proximus*) active before a decrease in influent phosphate.

However, under low-phosphate conditions, *A. regalis* (type I) remained the most active. While comparing previous studies (Fluorescence In Situ Hybridization) with the present study (metatranscriptomics) is challenging, the current study’s smooth conditioning cycles, conducted a week apart, likely influenced the results. Unlike previous studies that applied consecutive conditioning cycles until no phosphate was released, this study observed phosphate depletion 40 days after changing the influent. Although previous studies suggested that *Accumulibacter* type I were less efficient in adapting to a drastic change in phosphate conditions than type II, the present results indicate that they are capable of adapting to smooth perturbations. Furthermore, the presence and development of *Propionivibrio* during the same period may have affected the competition between the two types of *Accumulibacter*.

### 4.3 Multiple EPS pathways expressed by *Accumulibacter* and the flanking community

Genes from multiple EPS pathways were transcribed during the granulation process suggesting a wide variety of EPS. The complexity and the presence of multiple EPS in AGS as well as changes in the composition has been previously highlighted (Weissbrodt et al., 2013). The different EPS certainly play different roles in the formation and stability of the biofilm matrix. Although the EPS were not quantified in this present work, the metatranscriptomics allowed to identify the potential microbial EPS producers.

*Accumulibacter* transcribed genes from multiple EPS pathways in all time point and most of them confirmed results previously described. Dueholm et al. (2023) identified genes cluster of Poly-N-acetylglucosamine (PNAG) in *Accumulibacter* genomes, as well as Pel operon in *Propionivibrio*. PNAG are partially N-acetylated polysaccharides that can influence synthesis, translocation and adhesiveness depending on the microorganisms (Flemming et al., 2023).

In their study, Tomás-Martínez et al. (2021) identified genes related to the nonulosonic acids (NulOs) pathways in *Accumulibacter* genomes and in proteomics results. NulOs are nine-carbon monosaccharides primarily found in eukaryotes and pathogenic bacteria. Common types include neuraminic acids and ketodeoxynonulosonic acids, with legionaminic and pseudaminic acids specific to bacteria. De Graff et al. (2019) demonstrated that sialic acids protect the EPS matrix from enzymatic degradation by binding to terminal positions in carbohydrate chains. An inverse transcription trend between pseudaminic and legionaminic acids genes suggests a shift in EPS composition during granulation, potentially due to regulation systems or changes in *Accumulibacter* species composition. In their phylogenetic analysis to determine the diversity of NulOs possibly produced by *Accumulibacter*, Tomás-Martínez et al. (2021) showed that most of the *Accumulibacter* metagenome-assembled-genomes (MAGs) contain the NeuAc synthase gene but not all, possibly due to incomplete MAGs or true genetic absence. Additionally, the expression of neuraminic acid binding proteins, permease, and tripartite ATP-independent periplasmic transporter proteins indicates that *Accumulibacter* recycles NulOs to conserve energy (Tomás-Martinez et al., 2021). Similarly, *Propionivibrio, Azonexus*, and *Contendobacter* expressed genes for recycling neuraminic acids but not for NeuAc synthase, suggesting they reuse NulOs produced by *Accumulibacter*. Further genomic investigation could confirm the presence of the NeuAc gene, indicating potential but unused capability for neuraminic acid production. *Flavobacterium* transcribed some genes of the pseudaminic acids pathway. However, it is not so clear from the literature and the results if they are involved in granule formation and sialic acids production.

The metatranscriptomics results show that aside *Accumulibacter*, other microorganisms (*Rhodoferax, Propionivibrio, Contendobacter* and *Azonexus*) transcribed genes from the other EPS types, such as alginate which is an anionic linear polysaccharide composed of mannuronic acid and guluronic acid residues. Mature granules exhibit higher concentration of ALE in most of the studies and could be the results of the microbial compression during granule maturation (Zahra et al., 2022). A higher ALE content was usually associated to the presence of PAOs and GAOs such as *Accumulibacter* and *Propionivibrio* respectively (Zahra et al., 2022). Another type of EPS, Curli proteins or amyloid adhesins, were transcribed by *Kaistella*, mostly when the granules were bigger. These amyloids are β-sheet-rich proteins and have already been identified in activated sludge flocs and AGS (Christiaens et al., 2022; Lin et al., 2018). The production of amyloid has been suggested for ammonia-oxidizing bacteria, *Zoogloea*, and some GAO such as *Competibacter*. In this study, the activity of these microorganisms was relatively low and it could not be detected that they expressed amyloid adhesins genes.

The present metatranscriptomics analysis allowed to identify the microorganisms involved in the EPS production and suggests which type of EPS potentially composed the granule extracellular matrix. However, the analysis indicated that even when working with an enriched-biomass, other populations could be involved in the EPS formation and/or stability. In most of the studies, the role of these flanking populations is often under estimated. Additionally, the analyses were conducted as an overview of the microbial activity in the reactor during the granulation, but there could be heterogeneity at the granule level. Indeed, as shown by Leventhal et al. (2018), individual granules can be composed of a specific microbial community leading to potentially a different EPS composition.

Finally, all along the experiment the fast-settling granules remained. Neither the change of the influent composition, the modification of the microbial community nor the potential change in the EPS composition did cause a disruption of the granules. It could be because of the slow dynamics in these modifications and, the remaining presence and activity of *Accumulibacter*. Instead of a disruption, a modification in the granule shape from smooth to grainy was observed, following the change in the PAO to GAO metabolism and the associated microbial populations. The granule shape can be related with the predominant organisms such as described by Weissbrodt et al. (2013). However, it could also be a natural maturation of the granules (Mills et al., 2024) and further investigations correlating image analysis and microbial activity would be of great help to better understand and manage the granule life cycle.

## 5 Conclusions

Extracellular polymeric substances are central in the AGS technology by aggregating a diverse microbial community. In this study, *Accumulibacter* was the most active population of the microbial community during granulation and expressed genes of several extracellular polymeric substance pathways. Following a long period with low phosphate concentrations during which a GAO metabolism was established, *Accumulibacter* was still present and active alongside with the GAO *Propionivibrio. Propionivibrio* highly expressed genes involved in the EBPR process and EPS production. The pattern of expressed EPS pathways was different for this genus, but the granules were still large indicating that the change in microbial community composition had no major impact on granule stability. The role of the flanking community which included the PAO *Azonexus*, the GAO *Contendobacter* and *Competibacter, Flavobacterium, Rhodoferax, Kaistella*, and *Zoogloea*, should not be underestimated as they contributed to a considerable proportion of the transcriptome. Metatranscriptomics represented the ensemble of the reactor biomass at specific stages, but each granule can be composed of different microbial communities and therefore contain different EPS. To have a clearer view on the granulation process, further analyses should be conducted at granule level, by differentiating granules and linking their active microbial community members to the composition of EPS and the concentration of the quorum sensing molecules.

## 7 Author Contributions

**Laëtitia Cardona**: Conceptualisation, Formal analysis, Visualisation, Investigation, Writing - Original Draft. **Jaspreet Singh Saini**: Data curation, Writing – Review & Editing. **Pilar Natalia Rodilla Ramírez**: Writing – Review & Editing. **Aline Adler**: Writing – Review & Editing. **Christof Holliger**: Conceptualisation, Writing – Review & Editing, Supervision, Project Management, Funding acquisition.

## 9 Acknowledgments

We thank Emmanuelle Rohrbach (LBE, EPFL) for the molecular biology preparation work, Stéphane Marquis for the reactor monitoring and Marc Deront for the informatic support. Xenia Bender, Emylène Ostertag, Alyssa Etter and Alessandro Scapuso (LBE, EPFL) are also acknowledged for their help with the reactors monitoring and molecular biology preparations.

## 10 Declaration of Generative AI and AI-assisted technologies in the writing process

During the preparation of this work the author used Paperpal in order to improve and trim the text. After using this tool, the authors reviewed and edited the content as needed and take full responsibility for the content of the publication.

## Supplementary Material

**Appendix A. Supplementary figure**

Complementary figures: Biomass pictures and gene expression in aerobic phase

**Appendix B. Supplementary data**

Summary of the 16S rRNA gene amplicon sequencing

**Appendix C. Supplementary data**

Summary of the bioinformatic results, featureCount results

**Appendix D. Supplementary data**

Gene annotation

**Appendix E. Supplementary data**

List of genes of interest

## 12 Data Availability Statement

Raw metatranscriptomics reads and the de novo assembly after clustering using CD-HIT have been deposited at the Read Sequence Archive (SRA) and Transcriptome Shotgun Assembly (TSA) under the BioProject ID PRJNA1144857. The raw reads from the 16S rRNA gene amplicon sequencing are available under the BioProject ID PRJNA1125294.

