## Appendix A for "Multiple extracellular polymeric substances pathways expressed by *Accumulibacter* and the flanking community during aerobic granule formation and after influent modification"

EPFL - Ecole Polytechnique Federale de Lausanne

ENAC IIE LBE

CH B2 407

Station 6

1015 Lausanne - Switzerland

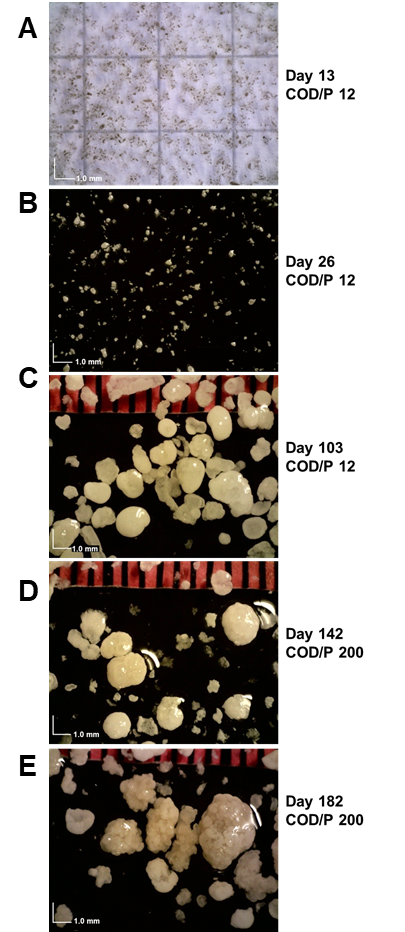

Supplementary figure 1. Evolution of the biomass before and after decreasing the phosphate concentration in influent composition to reach a COD/P ratio of 200. A) After 13 days of operation (COD/P 12). B) After 26 days of operation (COD/P 12). C) After 103 days of operation (COD/P 12). D) After 142 days of operation (COD/P 200). E) After 182 days of operation (COD/P 200)

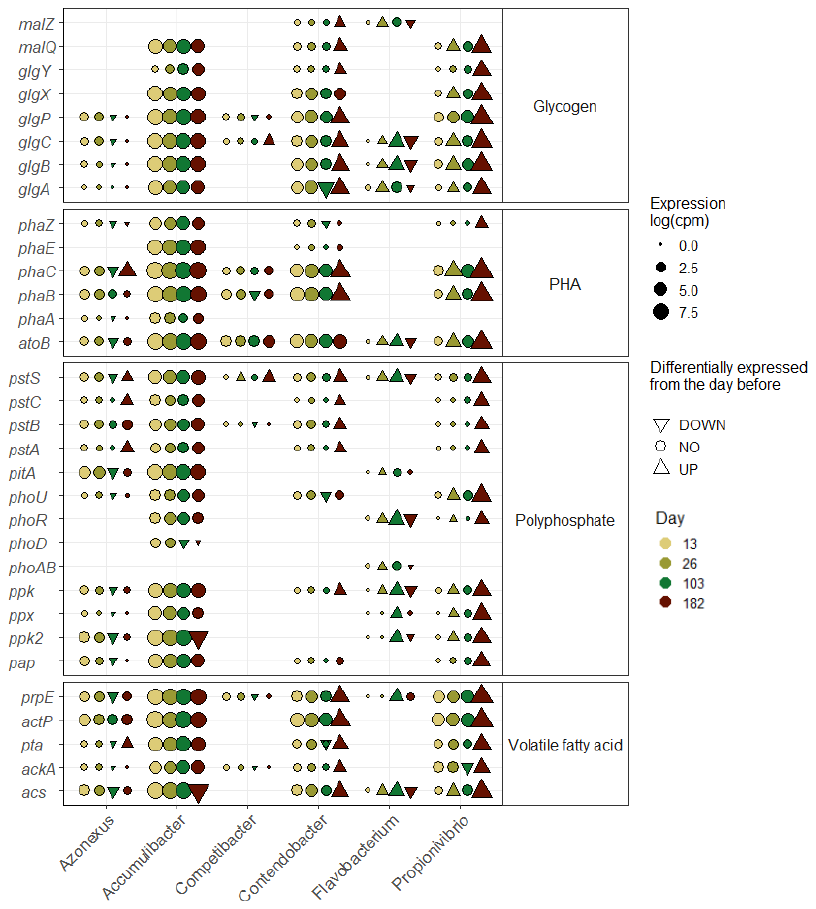

**Supplementary Figure 2. Expression of Enhanced Biological Phosphate Removal related genes at the aerobic phase.** Level of expression (log(cpm)) of genes per day for different genera. Differential gene expression analysis was done between two time points (26 versus 13, 103 versus 26 and 182 versus 103) and the significative differences (log-fold change > 2 and pvalue < 0.01) are represented by a triangle (up-pointing for **up regulation** and down-pointing triangle for **down-regulation**).

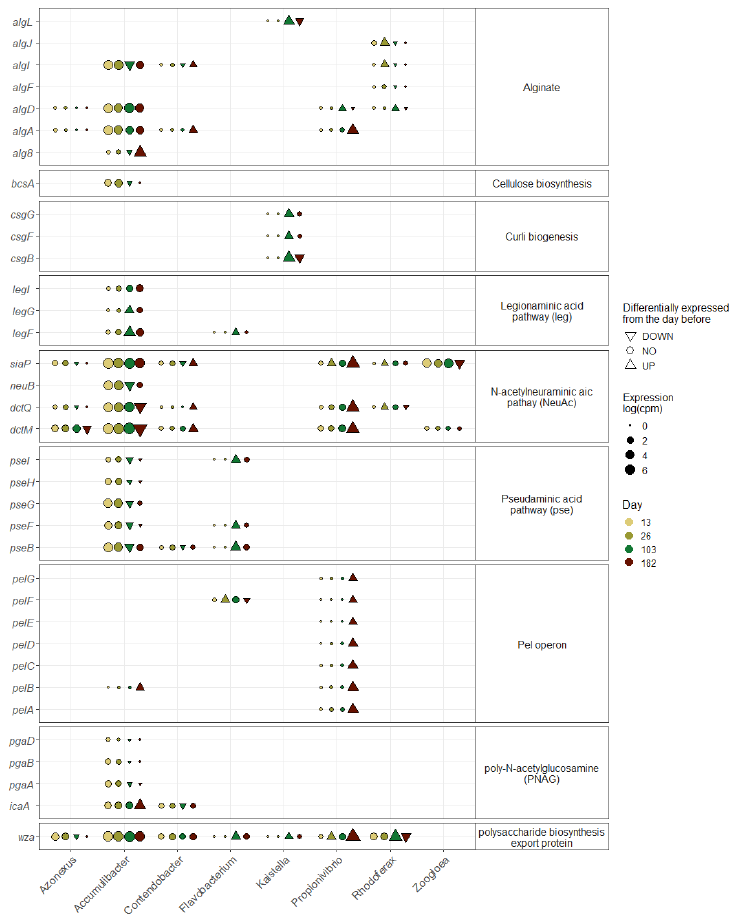

Supplementary Figure 3. Expression of biofilm related genes at the aerobic phase. Level of expression (log(cpm)) of genes per day for different genera. Differential gene expression analysis was done between two time points (26 versus 13, 103 versus 26 and 182 versus 103) and the significative differences (log-fold change > 2 and pvalue < 0.01) are represented by a triangle (up-pointing for up regulation and down-pointing triangle for down-regulation).
